# Effective prophylaxis of COVID-19 in rhesus macaques using a combination of two parentally-administered SARS-CoV-2 neutralizing antibodies

**DOI:** 10.1101/2021.05.26.445878

**Authors:** Brandon J. Beddingfield, Nicholas J. Maness, Alyssa C. Fears, Jay Rappaport, Pyone Pyone Aye, Kasi Russell-Lodrigue, Lara A. Doyle-Meyers, Robert V. Blair, Ann M. Carias, Patrick J. Madden, Ramon Lorenzo Redondo, HongMei Gao, David Montefiori, Thomas J. Hope, Chad J. Roy

## Abstract

SARS-CoV-2 is a respiratory borne pathogenic beta coronavirus that is the source of a worldwide pandemic and the cause of multiple pathologies in man. The rhesus macaque model of COVID-19 was utilized to test the added benefit of combinatory parenteral administration of two high-affinity anti-SARS-CoV-2 monoclonal antibodies (mAbs; C144-LS and C135-LS) expressly developed to neutralize the virus and modified to extend their pharmacokinetics. After completion of kinetics study of mAbs in the primate, combination treatment was administered prophylactically to mucosal viral challenge. Results showed near complete virus neutralization evidenced by no measurable titer in mucosal tissue swabs, muting of cytokine/chemokine response, and lack of any discernable pathologic sequalae. Blocking infection was a dose-related effect, cohorts receiving lower doses (6, 2 mg/kg) resulted in low grade viral infection in various mucosal sites compared to that of a fully protective dose (20 mg/kg). A subset of animals within this cohort whose infectious challenge was delayed 75 days later after mAb administration were still protected from disease. Results indicate this combination mAb effectively blocks development of COVID-19 in the rhesus disease model and accelerates the prospect of clinical studies with this effective antibody combination.

## Introduction

SARS-CoV-2 is a pathogenic beta-coronavirus that is the source of a current worldwide pandemic ^1^. The virus has proved to be of relatively low pathogenicity, and passes easily via respiratory droplet between the shedding infectious to naïve host. The combination of inability to readily kill immunocompetent hosts, but rather use the respiratory system as an efficient vectoring pathway to infect others has made the resultant disease (COVID-19) one of the most prolific in human history, infecting hundreds of millions, and killing >2.7 million worldwide ^2^. Infection results in a systemic but not bloodborne viral invasion, and associated clinical state (COVID-19) can be characterized in most as a mild to moderate upper respiratory, self-limiting disease ^3,4^. In some, however, including the aged, obese, and immunocompromised, the severe phenotype of the disease can result in significant pathologic sequalae and death^4–8^. This virus has been shown to be highly transmissible by numerous routes, including as an airborne respiratory pathogen. The human infectious dose is remarkably low, estimated as low as approximately 100 virions, and is one of the primary reasons that the virus is such a highly successful transmissible pathogen. Viral infection is mediated primarily through the angiotensin-converting enzyme 2 receptor (ACE2) and species with tissues rich in the ACE2 receptor (including man) primarily govern innate susceptibility to this virus^9–12^. The course of infection typically initiates in the upper mucosa, and quickly spreads to distal mucosal tissues, including the lower gastrointestinal tract by day 1-2 post infection. After a productive infection with associated respiratory clinical signs, viral titers recede in these same tissues and viral elimination takes place through a combination of innate response and short-lived neutralizing antibodies. Sequestration of virus in distal, ‘sanctuary’ tissues like the lower gastrointestinal tract, which is rich in ACE2 receptors clusters, has been reported clinically and in animal modeling studies that extend past a month of observation. There are few animal species that emulate the human condition and are consistent with the biological endpoints associated with COVID-19 and can be used as a disease model for evaluation^13–15^. The nonhuman primate, and specifically the rhesus macaque (*Macaca mulatta*) has emerged as a reasonable higher-order species from which advanced COVID-19 product evaluation studies can be performed. Experimental COVID-19 in the Rhesus is induced with a relatively large dose (1e+6 TCID) of SARS-CoV-2 delivered mucosally (intranasal (IN), intratracheal (IT)) that results in development of a clinically mild to moderate upper respiratory infection, with resolution 14-21 days after experimental infection. This disease model has been used extensively in pathogenesis, vaccine, and therapeutic evaluations^16–23^. The rhesus model is infected using sequential mucosal multiroute (IN, IT) exposure and was chosen as the species and route combination for the purposes of this evaluation.

Antibody prophylaxis is one of the hallmark strategies for prevention of infection by an infectious agent and this approach has been used for a number of viral pathogen, including SARS-CoV-2^24–30^. The antibodies evaluated in the macaque COVID model originated from COVID-19-convelescent donor blood^3,25,31^. Although both antibodies target the SARS-CoV-2 spike trimer, the viral protein that controls binding to ACE2 receptor, one (C135-LS) targets a highly conserved epitope outside the ACE2 binding site, and provides for non-overlapping epitope coverage when used in combination with another categorically distinct antibody (C144-LS) that binds to the receptor binding domain (RBD) in a more conventional manner. Both revealed to be highly potent in neutralizing SARS-CoV-2 after exhaustive structure resolution, affinity determination, and respective neutralizing capacity^25^. Both antibodies contained LS mutations at amino acids 428 and 434, which leads to a increased interaction with FcRn^27,32^. This preferential interaction with FcRn, which is a key regulator of plasma IgG homeostasis, increases the frequency of FcRn mediated rescue of antibodies from degradation in the lysosome. This preferential salvage increases the antibody PK. In the example of these 2 monoclonal antibodies, the LS mutation leads to a highly stable and long-lasting protection from COVID-19 infection.

In this study, we evaluated the C144-LS/C135-LS neutralizing antibody combination to prevent SARS-CoV-2 infection in rhesus macaques. Initially, the kinetics of C144-LS/C135-LS were measured in serum and bronchoalveolar lavage fluid (BAL) in uninfected macaques using a surrogate ACE2/RBD binding ELISA and pseudovirus neutralization assay. Both measures provided prevailing neutralization and RBD binding levels over time, which allowed for optimal timing of planned infectious challenge. In another cohort of macaques, groups of animals were administered three dosage levels of C144-LS/C135-LS and then challenged mucosally with SARS-CoV-2 three days later. In a corollary study, the animals dosed in the kinetics study were delay-challenged with SARS-CoV-2 75 days after receiving antibody administration.

## Materials and Methods

### Study Approval

The Tulane University Institutional Animal Care and Use Committee approved all procedures used during this study. The Tulane National Primate Research Center (TNPRC) is accredited by the Association for the Assessment and Accreditation of Laboratory Animal Care (AAALAC no. 000594). The U.S. National Institutes of Health (NIH) Office of Laboratory Animal Welfare number for TNPRC is A3071-01. Tulane University Institutional Biosafety Committee approved all procedures for work in, and removal of samples from, Biosafety Level 3 laboratories.

### Virus and Cells

Virus used for animal inoculation was strain SARS-CoV-2; 2019-nCoV/USA-WA1/2020 (BEI# NR-52281) prepared on subconfluent VeroE6 cells (ATCC# CRL-1586) and confirmed via sequencing. VeroE6 cells were used for live virus titration of biological samples and were maintained in DMEM (#11965092, Thermo Scientific, USA) with 10% FBS.

### Animals and Procedures

A total of 19 Rhesus macaques (*Macaca mulatta*), between 3 and 11 years old, were utilized for this study. All Rhesus macaques (RMs) were bred in captivity at TNPRC. Three RMs were infused with 20 mg/kg of an equal (10 mg/kg each) combination of C-135-LS and C-144-LS monoclonal antibodies raised against SARS-CoV-2 spike protein and monitored for RBD binding as well as neutralizing activity routinely for 75 days to determine pharmacokinetics. Sixteen of the RMs were infused with 20, 6, or 2 mg/kg mAb cocktail three days before challenge. They were then exposed via intratracheal/intranasal (IT/IN) installation of viral inoculum (1mL intratracheal, 500 μL per nare, total delivery 2e+6 TCID_50_). The three RMs used for pharmacokinetic studies were exposed similarly 75 days after initial infusion to assess responses to delayed challenge. Animal information, including time to challenge and mAb dosage, can be found in Table S1.

The animals were monitored twice daily for the duration of the study, with collections of mucosal swabs (nasal, pharyngeal, rectal) as well as fluids (bronchioalveolar lavage, cerebrospinal fluid, urine) were taken pre-exposure as well as post-exposure days 1, 3, and at necropsy. Blood was collected pre-exposure, as well as 1, 2, 3, 5 and at necropsy. Physical examinations were performed daily after exposure, and necropsy occurred between 7 and 9 days post-exposure. No animals met humane euthanasia endpoints during this study. During necropsy, tissues were collected in media, fresh frozen, or in fixative for later analysis.

### Quantification of Viral RNA in Swab and Tissue Samples

Viral load in tissues, swabs and BAL cells and supernatant was quantified using RT-qPCR targeting the nucleocapsid (genomic and subgenomic) or envelope gene (subgenomic) of SARS-CoV-2. RNA was isolated from non-tissue samples using a Zymo Quick RNA Viral Kit (#R1035, Zymo, USA) or Zymo Quick RNA Viral Kit (#D7003, Zymo, USA) for BAL cells, per manufacturer’s instructions. RNA was eluted in RNAse free water. During isolation, the swab was placed into the spin column in order to elute the entire contents of the swab in each extraction. BAL supernatant was extracted using 100 μL, and serum was extracted using 500uL. Viral RNA from tissues was extracted using a RNeasy Mini Kit (#74106, Qiagen, Germany) after homogenization in Trizol and phase separation with chloroform.

Isolated RNA was analyzed in a QuantStudio 6 (Thermo Scientific, USA) using TaqPath master mix (Thermo Scientific, USA) and appropriate primers/probes (Table S2) with the following program: 25°C for 2 minutes, 50°C for 15 minutes, 95°C for 2 minutes followed by 40 cycles of 95°C for 3 seconds and 60°C for 30 seconds. Signals were compared to a standard curve generated using *in vitro* transcribed RNA of each sequence diluted from 10^8^ down to 10 copies. Positive controls consisted of SARS-CoV-2 infected VeroE6 cell lysate. Viral copies per swab were calculated by multiplying mean copies per well by amount in the total swab extract, while viral copies in tissue were calculated per microgram of RNA extracted from each tissue.

### Quantification of Live Virus in Swab and BAL Samples

Median Tissue Culture Infectious Dose (TCID_50_) was used to quantify replication-competent virus in swabs and BAL supernatant. VeroE6 ells were plated in 48-well tissue culture treated plates to be subconfluent at time of assay. Cells were washed with serum free DMEM and virus from 50uL of sample was allowed to adsorb onto the cells for 1 hour at 37°C and 5% CO_2_. After adsorption, cells were overlayed with DMEM containing 2% FBS and 1% Anti/Anti (#15240062, Thermo Scientific, USA). Plates were incubated for 7-10 days before being observed for cytopathic effect (CPE). Any CPE observed relative to control wells was considered positive and used to calculate TCID_50_ by the Reed and Muench method.

### Pathology and Histopathology

Animals were humanely euthanized following terminal blood collection. The necropsy was performed routinely with collection of organs and tissues of interest in media, fresh frozen, and in fixative. The left and right lungs were imaged and then weighed individually. A postmortem bronchoalveolar lavage (BAL) was performed on the left lower lung lobe. Endoscopic bronchial brushes were used to sample the left and right mainstem bronchi. One section from each of the major left and right lung lobes (anterior, middle, and lower) sample fresh, and the remaining lung tissue was infused with fixative using a 50 mL syringe, and saved in fixative. Fixed tissues were processed routinely, embedded in paraffin and cut in 5 μm sections. Sections were stained routinely with hematoxylin and eosin or left unstained for later analysis via fluorescent immunohistochemistry.

Histopathologic lesions identified in tissues were scored semiquantitatively by the same pathologist that performed the necropsies. Lesions were scored based on severity as the lesions being absent (−), minimal (+), mild (++), moderate (+++), or severe (++++). For statistical analysis of histopathologic lesions, the four treatment cohorts were grouped together, and the two control cohorts were grouped together and non-parametric pairwise comparisons were performed.

Fluorescent immunohistochemistry was performed on 5um sections of Formalin-fixed, paraffin-embedded lung were incubated for 1 hour with the primary antibodies (SARS, Guinea Pig, (BEI, cat#NR-10361) diluted in NGS at a concentration of 1:1000). Secondary antibodies tagged with Alexa Fluor fluorochromes and diluted 1:1000 in NGS were incubated for 40 minutes. DAPI (4’,6-diamidino-2-phenylindole) was used to label the nuclei of each section. Slides were imaged with Zeiss Axio Scan Z.1 slide scanner.

### Detection of Neutralizing Antibodies in Serum

The ability of antibodies in serum to disrupt the binding of the receptor binding domain (RBD) of SARS-CoV-2 spike protein to Angiotensin Converting Enzyme (ACE2) was assessed via the Surrogate Virus Neutralization Test (GenScript# L00847) using the included kit protocol modified per the following: Serum samples were diluted from 1:10 to 1:21, 870 in order to determine an IC_50_ for RBD/ACE2 binding. Pseudovirus neutralization testing of matched serum was performed using a SARS-CoV-2.D614G spike-pseudotyped virus in 293/ACE2 cells, with neutralization assessed via reduction in luciferase activity^33,34^.

### Cytokine/Chemokine Analysis

Characterization of the immune response via bead-based determination of cytokine and chemokine concentrations was performed using a Life Technologies Cytokine 37-plex Non-Human Primate Panel for Luminex platform (#EPX370-40045-901, Thermo Scientific, USA) according to manufacturer instructions in BSL-2+ conditions. Samples were plated in duplicate utilizing included diluents and standards after clarification by centrifugation. Plates were read on a Bio-Plex 200 System (Bio-Rad Laboratories, USA). Heat maps of log_2_ fold change from raw fluorescence values were generated using the heatmap package in RStudio and normalized to baseline. Hierarchical clustering was unsupervised. Red was used for fold-change increase and blue was used for fold-change decrease.

### Clinical Chemistries

Analysis of blood chemistries was performed using a Sysmex XT-2000i analyzer for EDTA collected plasma, or an Olympus AU400 chemistry analyzer for serum.

### Statistical Analysis

Data analysis was performed using Graphpad Prism 9.0.2 software (Graphpad Software, USA). Comparison of PCR viral load between groups was performed using a Kruskal-Walis ANOVA using the area under the curve of each individual combined into the dosage groups as compared to the control group. For comparison of TCID_50_ and day-by-day comparison of PCR viral load between groups, a Mann-Whitney t-test was used to compare between each dosage group and controls. Relationships between serum neutralization and viral loads were assessed via Spearman correlation. For histopathological scoring, comparisons were made via Mann-Whitney t-test. A two-way ANOVA with Tukey multiple comparisons test was utilized for cytokine/chemokine comparisons.

## Results

### Antibody Kinetics

A cocktail of C-135 and C-144 antibodies was administered to three Rhesus macaques in order to monitor levels of those antibodies over time (Fig. 1A). Antibody levels in serum, as assessed via competitive RBD/ACE2 binding ELISA, persisted beyond initial administration, with levels detectable day 75 post administration remaining to 10% of initial values in these long-term animals. (Fig. 1B). Serum antibody levels assessed via pseudovirus neutralization assay were similarly long-lived, with 75-day post administration levels up to 14% of their initial values (Fig. 1C). BAL levels of antibody were highest at day 3 post administration, resulting in subsequent challenge on day 3 post administration in the initial challenge groups (Fig. 1D).

**Figure 1.**
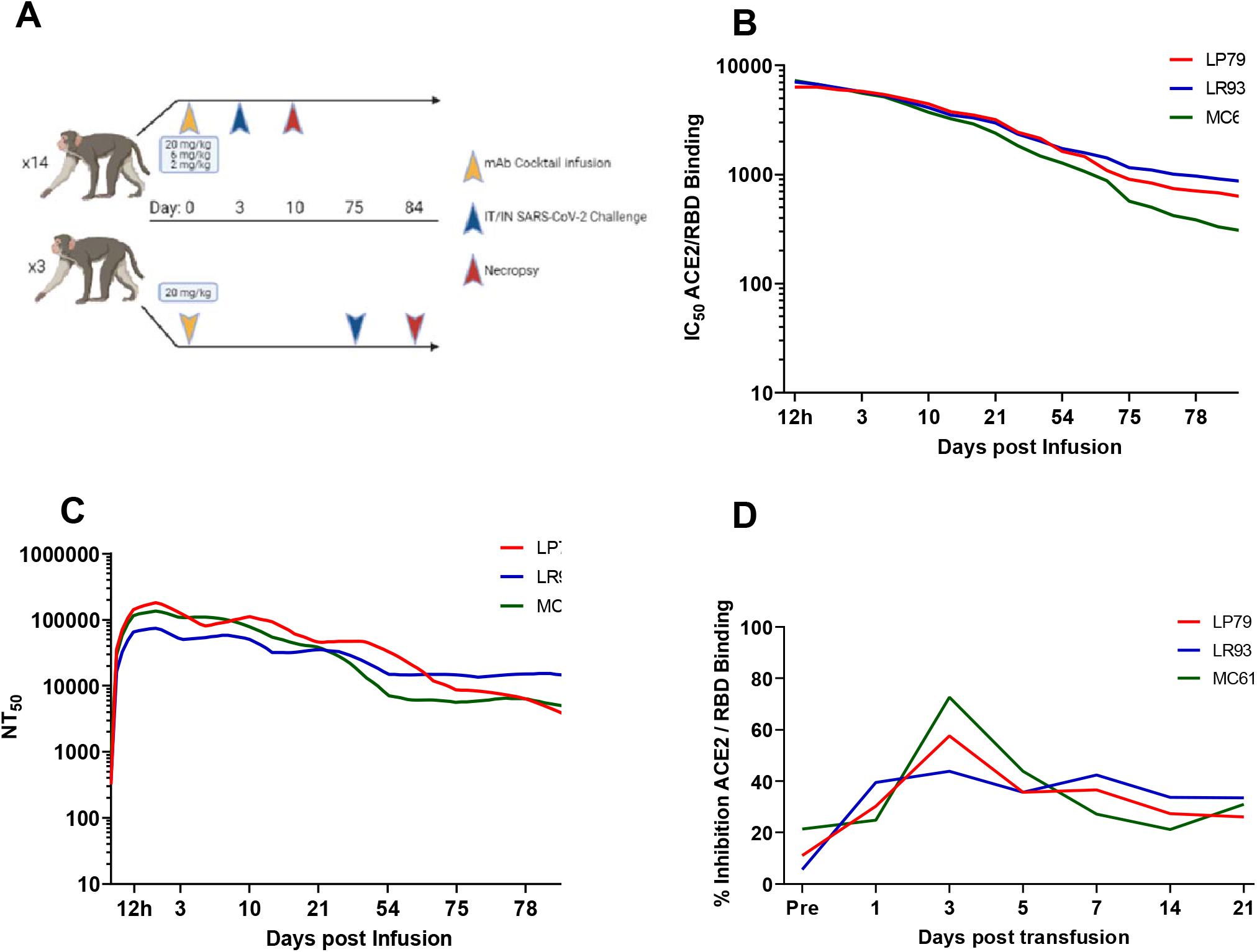
Study Design and Antibody Kinetics. A) Three different dosages of mAb cocktail consisting of equal amounts of C-135 and C-144 were administered three days before challenge. Separately, animals administered 20 mg/kg of mAb cocktail were challenged 75 days post administration. Antibody kinetics in serum were followed by both surrogate ELISA (B) and pseudovirus neutralization assay (C), and BAL supernatant antibody kinetics followed by surrogate ELISA (D).

### Protective Efficacy of mAbs After SARS-CoV-2 Challenge

Rhesus macaques were challenged three days post administration of monoclonal antibody cocktail, consisting of 10 mg/kg each of C-135 and C-144 and followed for seven days with regular collection of mucosal swabs. A separate cohort of animals from the initial kinetics experiment were challenged with SARS-CoV-2 by the same modality at 75 days post mAb administration. Viral load was monitored by qPCR analysis of the N gene genomic and the N and E gene subgenomic content as well as the TCID_50_ content, of the swabs and BAL cells (supernatant for TCID_50_). Viral loads were significantly lower in the 20 and 6 mg/kg groups than the control in all three PCR analyses in the pharyngeal swab (Fig. 2 A, E, I). The 20 and 6 mg/kg groups were significantly lower in the nasal swab for subgenomic N (Fig. 2F), with those in addition to the 2 mg/kg and 75 day challenge being significantly lower in the subgenomic E analysis (Fig. 2J). The 75 day challenge group had significantly lower bronchial brush viral loads than the controls in the genomic N and subgenomic E analyses (Fig. 2C and 2G, respectively). In the BAL cells, the 20, 6 and 2 mg/kg groups were significantly lower in genomic N content than controls (Fig. 2D), with all treatment groups being significantly lower than controls for subgenomic E (Fig. 2L), and the 75 day challenge group being significantly lower than controls for subgenomic N content (Fig. 2H). Exact p values can be found in Figure S3. All non-75 day challenge groups, when compared by day, had significantly lower viral loads than controls, the only exceptions being bronchial brush assayed for subgenomic E content (Table S4, Fig. S1). Tissue viral load analyses revealed lower viral loads in most respiratory tissues in treatment groups (Fig. S2 A-C). Non-respiratory tissues, predominantly digestive, from the low dose (2 mg/kg) group possessed higher viral loads than controls, which was driven by one individual (Fig. S2 D-F). Live virus, as measured by TCID_50_, was never detected in the swabs or BAL supernatant of the initial (Fig. 3 E-P) or delayed (Fig. 3 Q-T) treatment groups, despite detection in the control groups of each sample type (Fig. 3 A-D).

**Figure 2.**
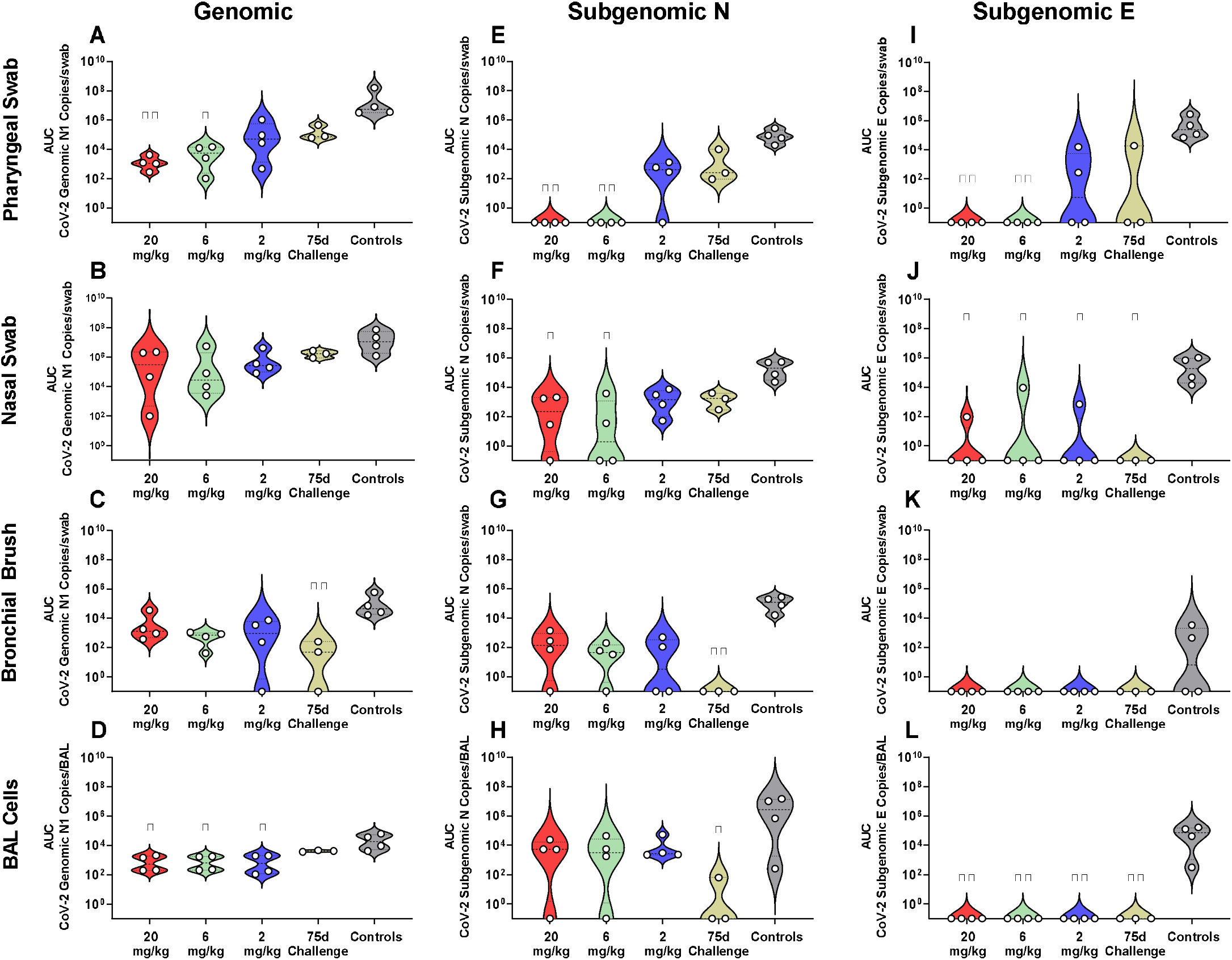
Viral Loads Assessed via Genomic and Subgenomic RT-qPCR During Challenge. Viral loads from pharyngeal swabs (A, E, I), nasal swabs (B, F, J), bronchial brushes (C, G, K) and BAL cells (D, H, L) assessed by RT-qPCR for genomic N (A-D), subgenomic N (E-H), and subgenomic E (I-L) content. Data are represented as area under the curve of each dosage group for each anatomical site and assay type. Groups were compared via Kruskal-Walis test comparing each dosage group to the control group. Asterisks represent significant difference from controls (*, p<0.05; **, p<0.01).

**Figure 3.**
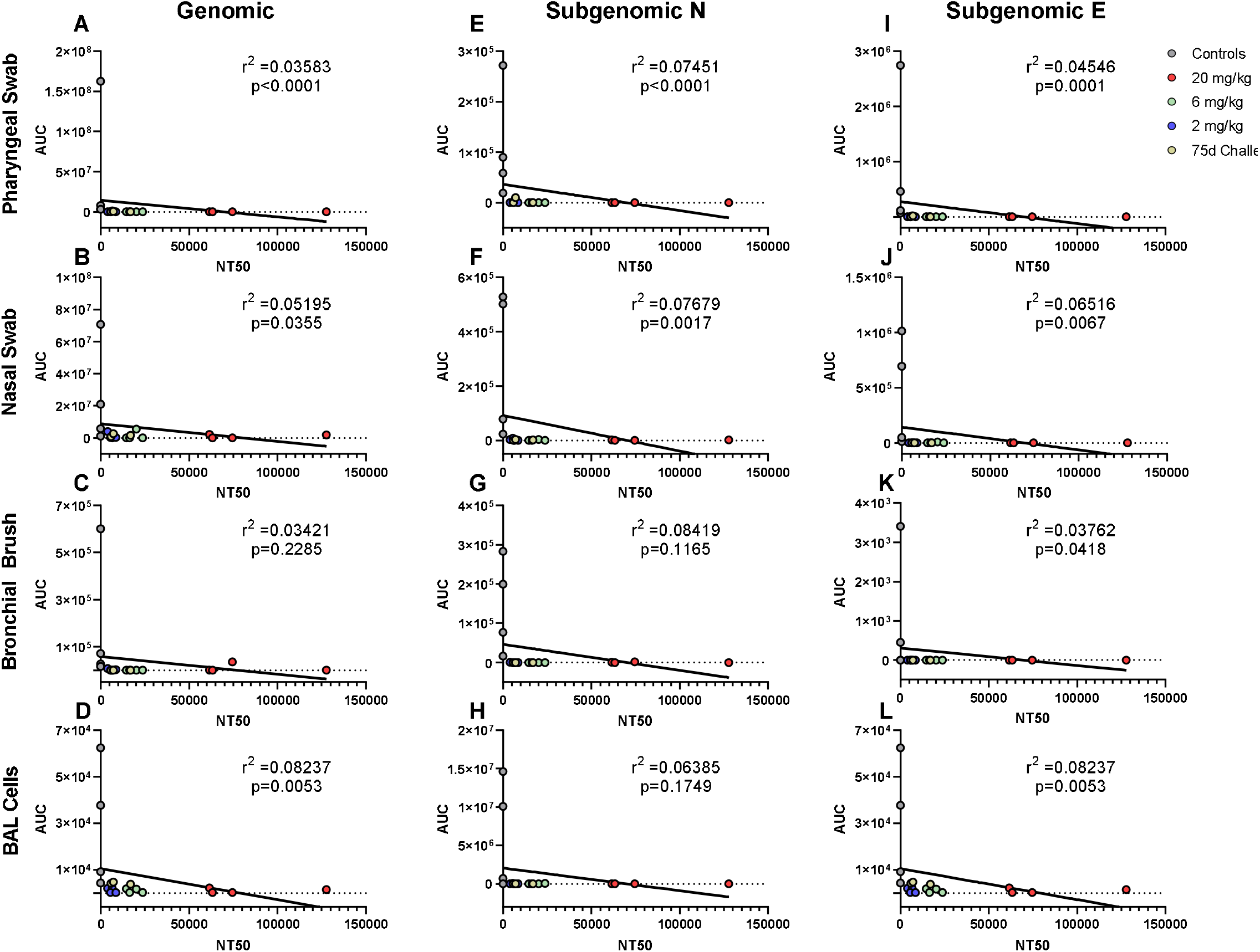
Correlations between Serum Antibodies and Viral Loads. Viral loads were correlated with neutralization capacity of serum collected 1 day post challenge from pharyngeal swabs (A, E, I), nasal swabs (B, F, J), bronchial brushes (C, G, K) and BAL cells (D, H, L) assessed by RT-qPCR for genomic N (A-D), subgenomic N (E-H), and subgenomic E (I-L) content. The NT_50_ of each was calculated via pseudovirus neutralization assay presented as reciprocal serum dilution. Nonparametric correlation with linear regression performed to determine the r^2^ value for each comparison.

### Antibody/Viral Load Correlations

Antibody levels of all groups, as assessed by pseudovirus neutralization, were inversely correlated to genomic N viral loads in pharyngeal, nasal and BAL cell samples (Fig. 4A, 4C, 4D, respectively). For subgenomic N content, pharyngeal and nasal were inversely correlated (Fig. 4E, 4F, respectively). All sites were inversely correlated for the subgenomic E assay (Fig. 4 I-L). When analyzed separately, antibody concentration inversely correlated with viral load in the 75 day challenge group in the bronchial brush and BAL cells for subgenomic N (Fig. 5G and 5H, respectively), as well as pharyngeal and nasal swabs and BAL cells for subgenomic E (Fig. 5I, 5J and 5L, respectively).

**Figure 4.**
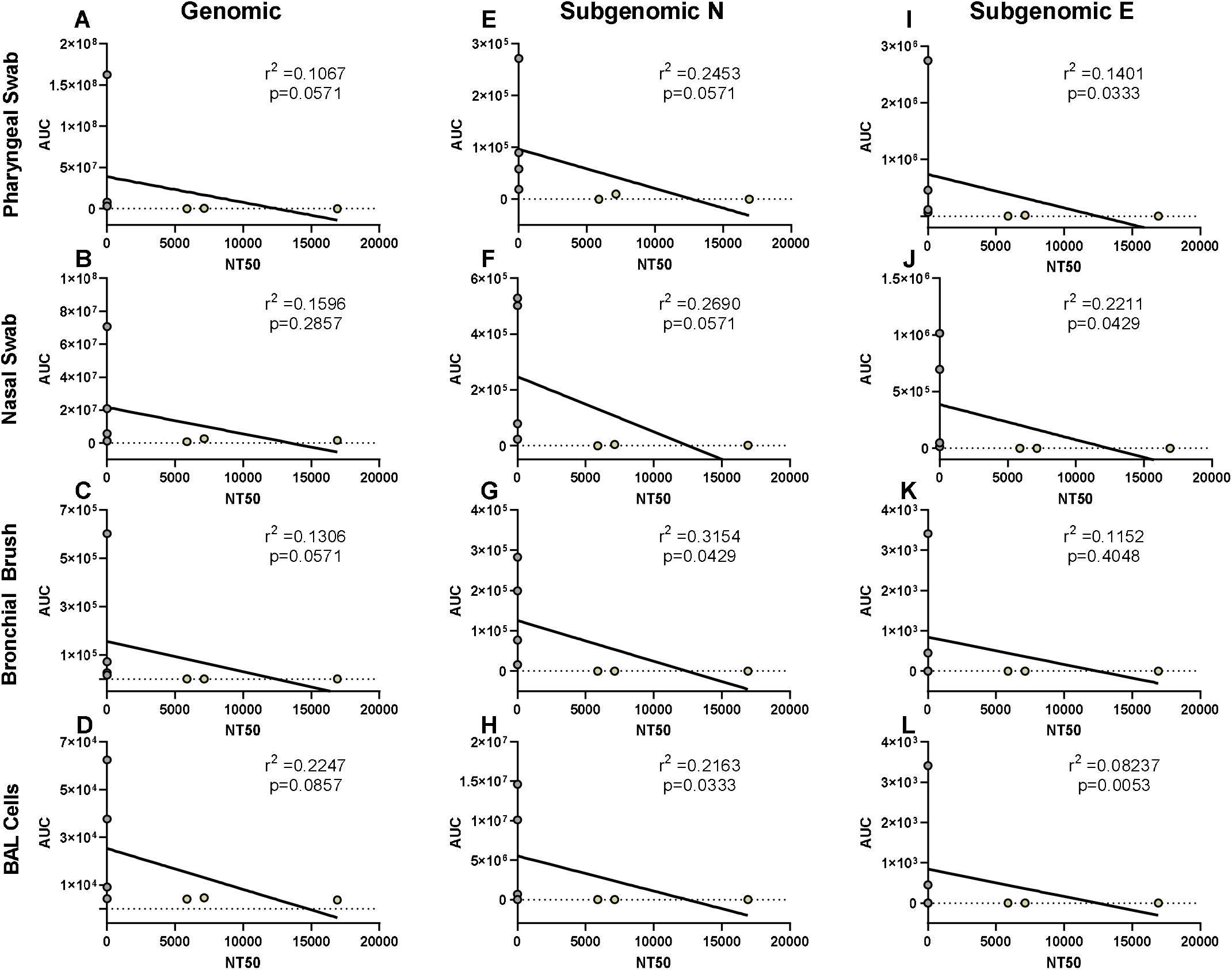
Correlations between Serum Antibodies of 75 Day Challenge Animals and Viral Loads. Viral loads were correlated with neutralization capacity of serum collected one day post challenge from pharyngeal swabs (A, E, I), nasal swabs (B, F, J), bronchial brushes (C, G, K) and BAL cells (D, H, L) assessed by RT-qPCR for genomic N (A-D), subgenomic N (E-H), and subgenomic E (I-L) content from the 75 day challenge group. The NT_50_ of each was calculated via pseudovirus neutralization assay presented as reciprocal serum dilution. Nonparametric correlation with linear regression performed to determine the r^2^ value for each comparison.

**Figure 5.**
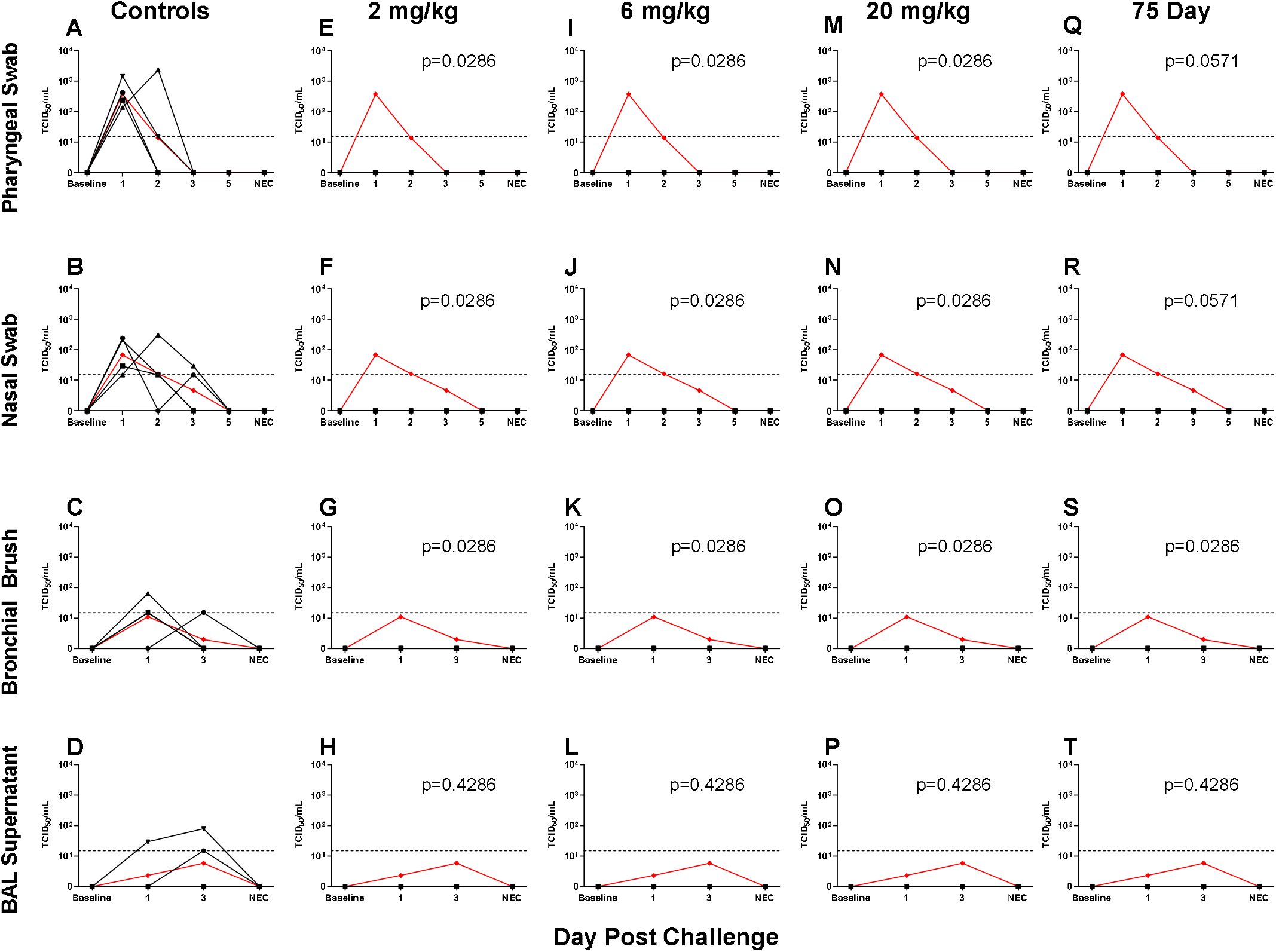
Viral Loads Assessed via Median Tissue Culture Infectious Dose (TCID_50_) During Challenge. Viral loads from pharyngeal swabs (A, E, I, M, Q), nasal swabs (B, F, J, N, R), bronchial brushes (C, G, K, O, S) and BAL supernatant (D, H, L, P, T) assessed TCID_50_ for controls (A-D), 2 mg/kg (E-H), 6 mg/kg (I-L), 20 mg/kg (M-P) and 75 day challenge (Q-T) groups. Data are represented as viral load calculated by the Reed and Muench method, with the red line indicating group mean and the dotted line indicating the quantification limit of assay.

### Lung Pathology

Gross pathology pathologic changes were noted in 6/19 RMs including 3/4 control animals and one animal each from the 2 mg/kg, 6 mg/kg, and delayed challenge groups. Gross lesions included small, multifocal tan areas of consolidation and pleuritis. Bronchial lymph node hyperplasia was also observed in the 3 control animals with pneumonia, but none of the treated animals (Fig. 6). Microscopically, interstitial pneumonia was seen in 14/15 treated RMs (only absent in one delayed challenge animal) and 4/4 control animals. In treated animals, interstitial pneumonia ranged from absent to mild and was significantly reduced compared to control animals which ranged from mild to severe (p=0.04). A significant reduction in the frequency and severity of type II pneumocyte hyperplasia was also observed between treated (4/15) and control (3/4) animals (p=0.04) (Table S5). No significant lesions were noted in other tissues. The myocarditis, nephritis, and gastrointestinal inflammation in the treated animals were interpreted as background lesions, and unlikely to be associated with therapy or SARS-CoV-2 infection.

**Figure 6.**
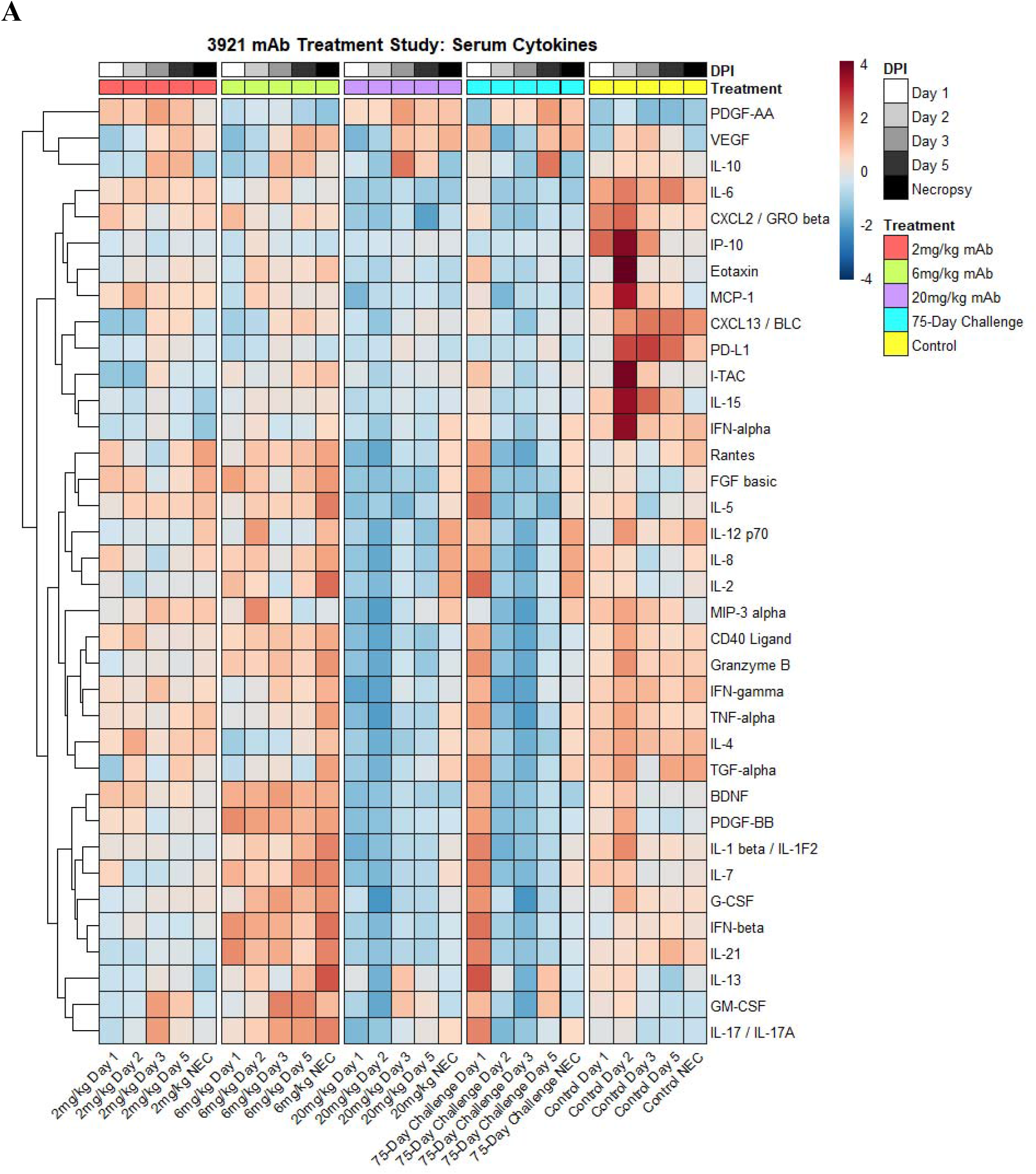
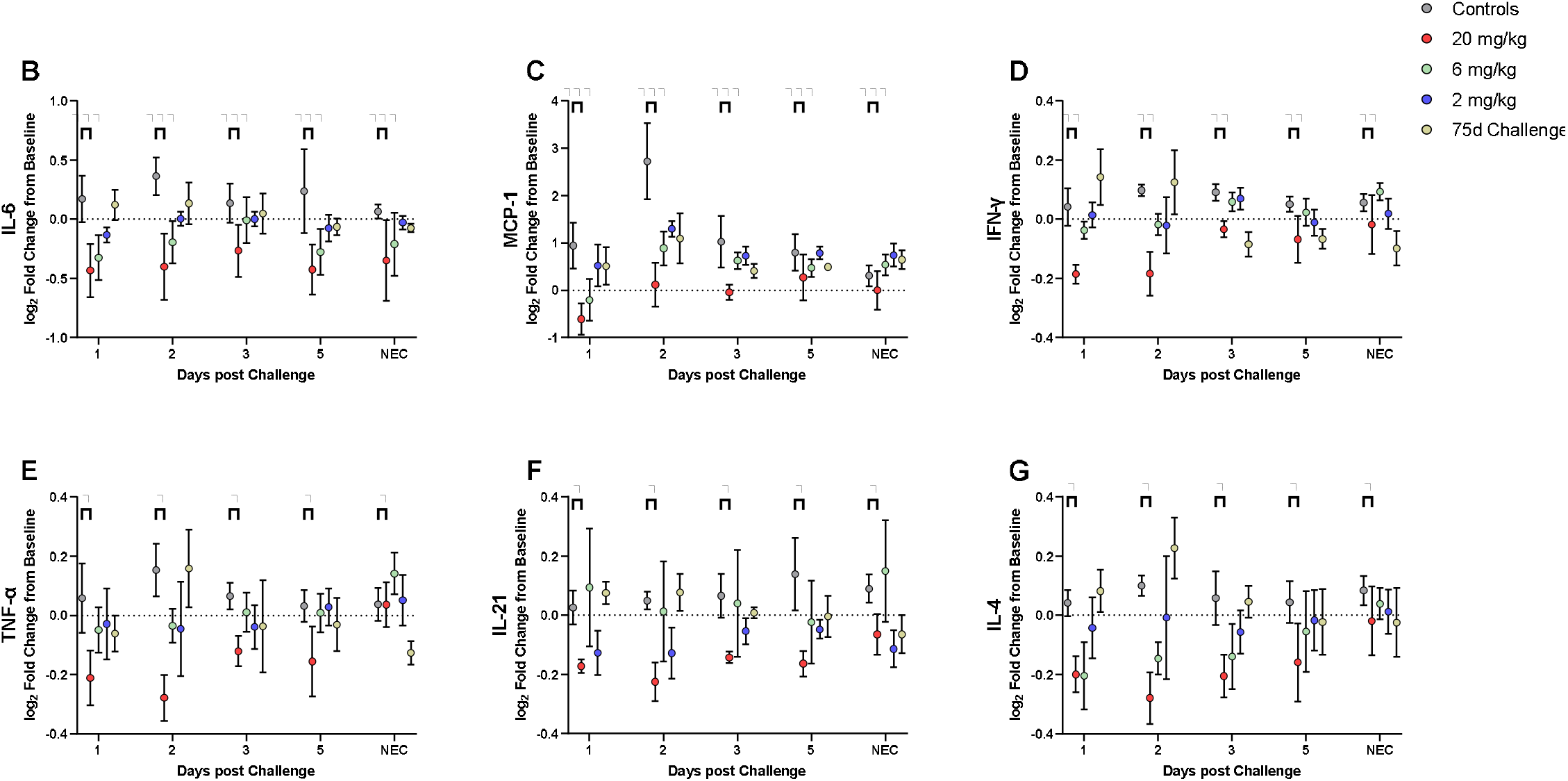
Cytokine and Chemokine Expression During Challenge. Cytokines were assessed during challenge for the different dosage groups represented as A) heat map of dosage groups vs. controls, and B-G) log_2_ fold change between dosage groups, with comparisons made via two-way ANOVA with Tukey’s multiple comparisons test (*, p<0.05; **, p<0.01; ***, p<0.001).

Fluorescent immunohistochemistry for SARS-CoV-2 was performed on one section of lung from each animal. Virus was detected in the lung of 2/4 control animals and 1/15 treated animals (LT54, 2 mg/kg). The lack of viral staining in two of the control animals may represent a sampling artifact, as virus was detected in BAL cells in all 4 controls (fig.7). This difference in viral staining between control and treatment groups trended toward but did not achieve statistical significance. (p=0.06; MW).

**Figure 7.**
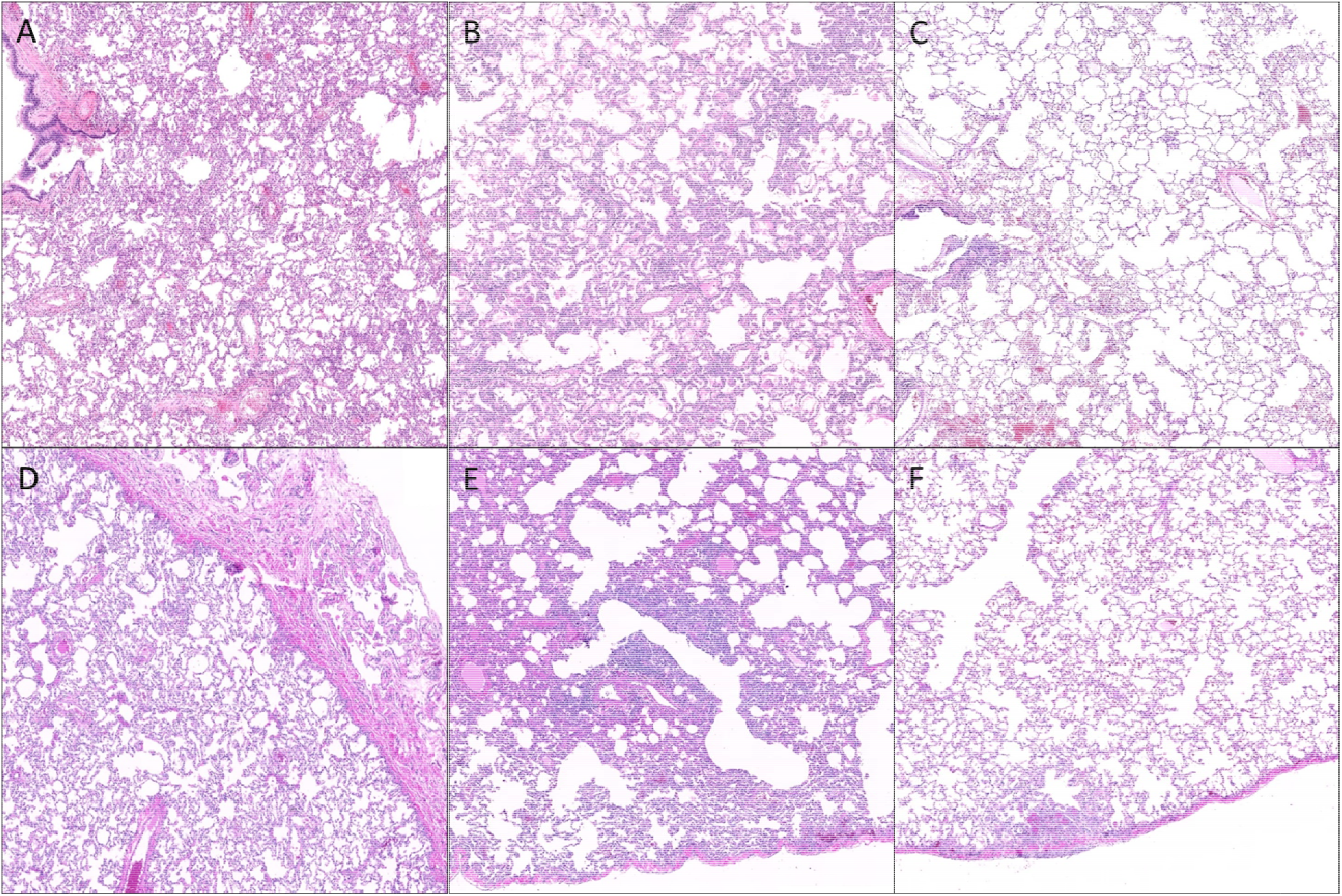
Histopathologic findings in the lungs of rhesus macaques challenged with SARS-CoV-2 following prophylactic administration of combination mAb therapy. A) Naïve (KF89). There is severe, widespread, interstitial inflammation. Inset: The inflammatory infiltrate is predominantly histiocytic with rare multinucleated cells (arrow). B) mAb control (IK92). There is severe, widespread interstitial inflammation. Inset: Alveolar septa are lined by type II pneumocytes that occasionally exhibit atypia (arrow). C) Delayed challenge (MC61). There is minimal interstitial inflammation with scattered aggregates of inflammatory cells (arrow) and hemorrhage (arrowhead). Inset: Inflammatory cells are predominately histiocytes (arrow). D) 2 mg/kg (MG10). There is mild interstitial inflammation (arrow) and pleuritis (asterisks). Inset: The interstitial inflammation is perivascular and composed of low numbers of lymphocytes and histiocytes. E) 6 mg/kg (ME55). There are multifocal regions of bronchointerstitial inflammation (arrows). Inset: Within areas of inflammation there is type II pneumocyte hyperplasia (arrow) and histiocytic infiltrate (arrowhead). F) 20 mg/kg (LR09). There are rare localized areas of subpleural inflammation (arrow). Inset: Within theses areas there is type II pneumocyte hyperplasia (arrow) and fibrin (asterisks). Bar = 0.5 mm.

### Cytokines/Chemokines

Cytokines were examined via a 36-plex luminex platform assay for all dosage groups during challenge with SARS-CoV-2. The 20 mg/kg dosage group exhibited blunted responses for pro-inflammatory cytokines (IL-6, TNF-α, IFN-α, IFN-γ, IL-12, IL-15, IL-21), chemokines (CXCL13, Eotaxin, IP-10, MCP-1, CXCL2), CD40L, PDGF-AA, PD-L1 and Granzyme B. Multiple differences were seen in the other dosage groups for IFN-α, IL-15, CD40L, PD-L1, CXCL13, and IP-10 (Fig. 8, Fig. S3).

**Figure 8.**
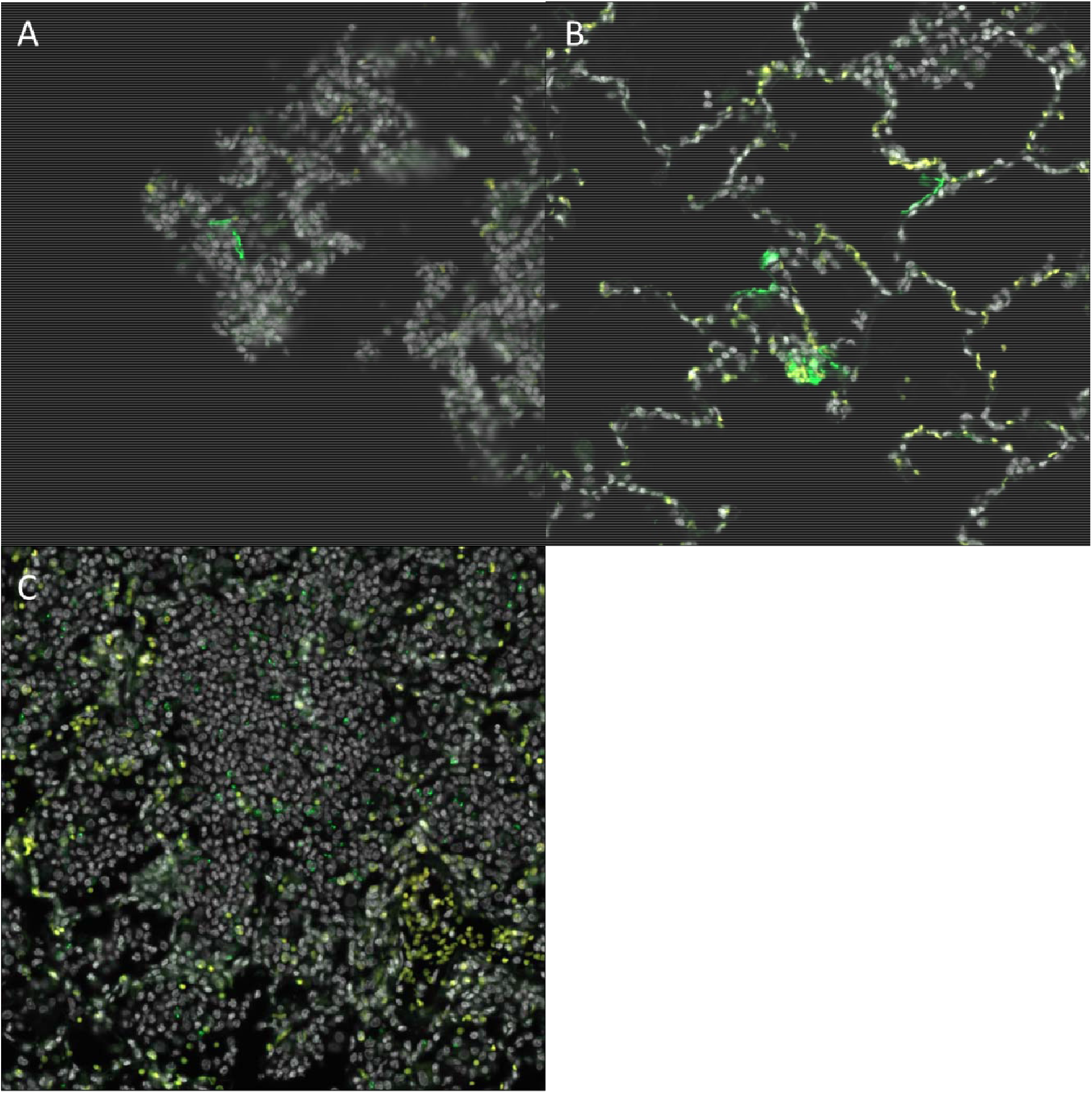
Fluorescent immunohistochemistry for SARS-CoV-2 in rhesus macaques following prophylactic treatment with combination mAb therapy. A) Naïve (LM30). Rare SARS-CoV-2 positive cells (green, arrows) are present lining alveolar septa. B) mAb control (IK92). SARS-CoV-2 positive cells are scattered throughout the lungs, often lining alveolar septa. C) 2 mg/kg (LT54). Within aggregates of inflammatory cells, there is scattered, punctate staining within inflammatory cells. Insets show a higher magnification of the SARS-CoV-2 positive signal described. DAPI = white, SARS-CoV-2 = green, Red = empty/autofluorescence. Bar =100 μm.

### Clinical Chemistries and Hematology

Hematological and chemistry values remained within the normal range throughout the study for most animals. Of note, the 75 day challenge group displayed lowered value changes in Alanine Aminotransferase (ALT) than the lowest (2 mg/kg) dosage group (Fig. S4); however, the values for both groups did not climb above normal range.

## Discussion

SARS-CoV-2 is a recently emerged and highly transmissible coronavirus. It is the etiologic agent for COVID-19 disease, and the source of an ongoing worldwide pandemic ^35^. More than two and a half million people have died worldwide from COVID-19 to date, with more than half a million deaths in the United States alone (Johns Hopkins University, 2021). Infection is associated with highly heterogenous disease sequelae, ranging from completely asymptomatic to severe acute respiratory distress and death. A number of vaccines are now becoming widely available that have remarkable potential to blunt the current pandemic. Passive infusion of highly potent human monoclonal antibodies (mAbs) that target the viral spike protein is another powerful tool with both prophylactic and therapeutic potential. Here, we tested the prophylactic potential of a combination of two mAbs administered either three or 75 days prior to SARS-CoV-2 respiratory challenge in rhesus macaques. We demonstrate that administration of these mAbs at a range of doses achieved near-complete protection from viral replication and disease even when challenge occurred 75 days after mAb administration. These data underscore the prospect for clinical use of these mAbs for infection prevention.

A number of animal models are now available for testing vaccines and therapeutics against SARS-CoV-2. Transgenic mice that express human ACE2 (hACE2), the receptor for SARS-CoV-2, offer a model of uniformly severe disease but data from this model requires cautious interpretation since it is likely that hACE2 is expressed in a variety of irrelevant cell types in this model. Siberian hamsters are likewise a model of moderate to severe disease and offer potential for testing novel therapeutics. Both of these models can be used for initial testing of therapeutic agents, but many of those agents will require testing in nonhuman primate models to assess translational potential before use in the clinic. The rhesus macaque model of COVID-19 recapitulates the mild to moderate disease shown in a large fraction of SARS-CoV-2 infected humans, but does not recapitulate the most extreme forms of the disease, including death. However, infection of macaques produces robust and quantifiable viral replication in respiratory and gastrointestinal tracts, as well as elevated pro-inflammatory cytokines, and acute virus-induced lesions in lungs ^23,36,37^, which can be compared between treated and untreated animals. Thus, rhesus macaques are a robust model for testing the efficacy of prophylactic administration of mAbs, particularly those that have shown potential in smaller animal models.

In this study we used a combination of two mAbs (C135-LS and C144-LS) with well-defined neutralization profiles that target defined epitopes in the viral spike protein ^27,38,39^. Both were engineered with LS mutations to increase half-life ^27,32^. The LS modifications are key in the potency of this combination of two monoclonals to provide potent protection 75 days after injection. We tested a range of doses (from 2 to 20 mg/kg) in our cohort of macaques challenged at 3 days post administration. A dose response was noted, as measured by the amount of viral subgenomic RNA expression, a correlate of active viral replication, in upper respiratory sites. However, even in the 2mg/kg group, viral replication after challenge was substantially lower than untreated control animals and no live virus was recovered from respiratory sites in any of our treated animals, suggesting a profound degree of protection, even at very low prophylactic doses.

An emerging aspect of the COVID-19 pandemic of critical importance is the emergence of viral variants, some of which appear to be bona fide escape mutations from either natural or vaccine induced humoral immunity ^40–43^. Importantly, although both C135-LS and C144-LS both target epitopes in the receptor binding domain (RBD) of spike, these epitopes are non-overlapping. Use of combinations of mAbs that target non-overlapping epitopes can prevent viral escape ^44,45^, which is particularly critical for SARS-CoV-2 given the recent emergence of several variants, including those that originated in South Africa (termed B.1.351) and Brazil (termed P.1), both of which include variants in spike that enable escape from host humoral immunity. It is particularly noteworthy that C144-LS targets an epitope that contains spike position 484, and neutralization provided by this mAb is nearly completely abrogated by the E484K variant ^39,45^, which is present in both B.1.351 and P.1. However, evidence to date suggest that C135-LS maintains full potency against these variants. Thus, in combination, C135-LS and C144-LS may offer broad protection against known circulating variants of SARS-CoV-2. Thus, this particular combination of mAbs has strong potential to protect individuals from infection and disease, even in the context of circulating variants that harbor escape mutations.

Taken together, our data strongly suggest that prophylactic use of combinations of mAbs, such as those we tested, administered systemically can provide durable and potent protection from mucosal infection with SARS-CoV-2. Indeed, this particular mAb cocktail is in clinical trial to determine real-world efficacy (clinicaltrials.gov identifier: NCT04700163). This approach could enable protection from disease for individuals that either cannot receive the available vaccines for any reason, or that work in high-risk environments and require enhanced protection from the virus.

**Table 1.**
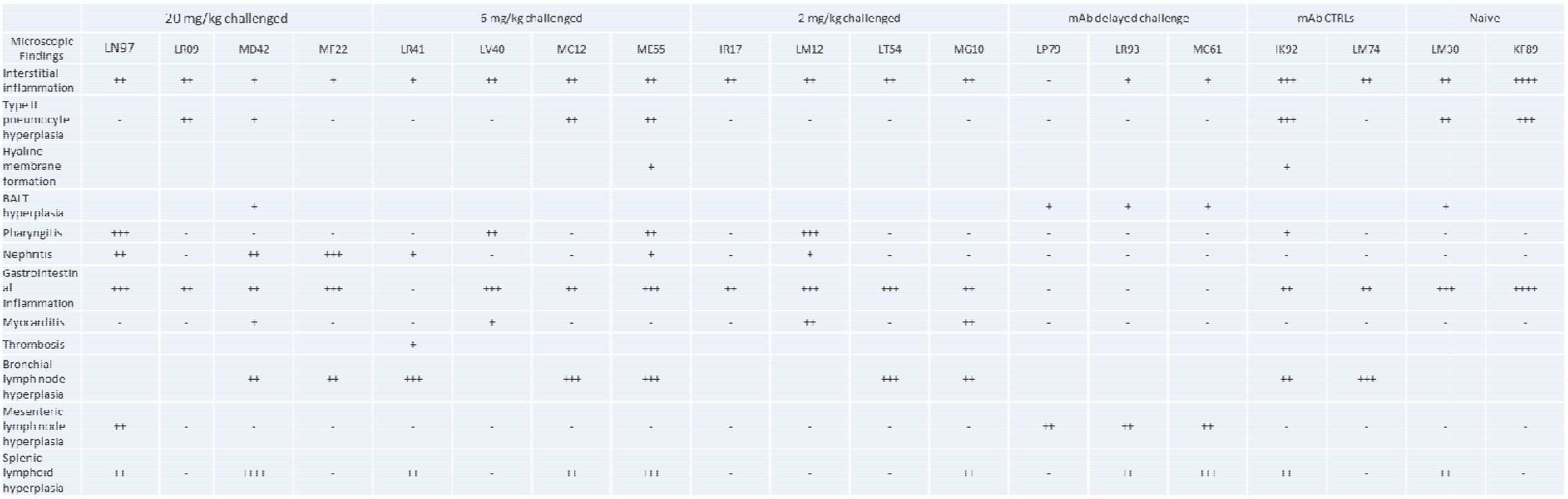
Histopathologic lesions in rhesus macaques receiving combined mAb therapy at dosages of 20, 6, and 2 mg/kg and challenged with SARS-CoV-2 compared to animals with delayed (75 days) challenge, those receiving a non-specific mAb, and naïve animals.

## Acknowledgements

We would like to thank Dr. Michel Nussenzweig for provision of the critical reagent monoclonal antibodies for this evaluation. We also acknowledge and thank Drs. Clint Florence, Jean Patterson, Que Dang, and Nancy Miller all at NIH/NIAID for invaluable scientific input on the study, and critical review of the manuscript. We thank Ms. Nadia Golden, Breeana Picou, Skye Spencer, and Krystal Hensley all members of the TNPRC High Containment Performance Core for their work We thank Angela Birnbaum for reviewing and optimizing all technical SOPs and overseeing the safety of this study.

