## Supplemental information for "Effective prophylaxis of COVID-19 in rhesus macaques using a combination of two parentally-administered SARS-CoV-2 neutralizing antibodies"

**Supplementary Figures**

| **Target** | **Primer/Probe Designation** | **Sequence** |
| --- | --- | --- |
| Genomic Nucleocapsid | 2019-nCoV_N1-F | 5’-GAC CCC AAA ATC AGC GAA AT-3’ |
|  | 2019-nCoV_N1-R | 5’-TCT GGT TAC TGC CAG TTG AAT CTG-3’ |
|  | 2019-nCoV_N1-P | 5’-FAM-ACC CCG CAT TAC GTT TGG TGG ACC-BHQ-3’ |
| Subgenomic Envelope | SgE-F | 5’-CGA TCT CTT GTA GAT CTG TTC TC-3’ |
|  | SgE-R | 5’-T GTG TGC GTA CTG CTG CAA TAT-3’ |
|  | SgE-P | 5’-FAM-ACA CTA GCC ATC CTT ACT GCG CTT CG-BHQ-3’ |
| Subgenomic Nucleocapsid | SgN-F | 5’-CGA TCT CTT GTA GAT CTG TTC TC-3’ |
|  | SgN-R | 5’-GGT GAA CCA AGA CGC AGT AT-3’ |
|  | SgN-P | 5’-56-FAM/TAA CCA GAA/ZEN/TGG AGA ACG CAG TGG G/3IABkFQ/-3’ |

**Table S1. Primer and Probe Sequences Utilized During this Study**

| Animal | Species | Age (years) | Source | Sex | Weight (kg) | Viral Dose (TCID_50_) | Exposure Route | mAb Dose (mg/kg) | Challenged post Infusion (Days) |
| --- | --- | --- | --- | --- | --- | --- | --- | --- | --- |
| LM74 | *Macaca mulatta* | 4 | TNPRC | Male | 6.2 | 2.0 x 10^6^ | IT/IN | 0 | 3 |
| IK92 | *Macaca mulatta* | 11 | TNPRC | Male | 6.9 | 2.0 x 10^6^ | IT/IN | 0 | 3 |
| KF89 | *Macaca mulatta* | 8 | TNPRC | Male | 8.2 | 2.0 x 10^6^ | IT/IN | 0 | 3 |
| LM30 | *Macaca mulatta* | 4 | TNPRC | Male | 8.1 | 2.0 x 10^6^ | IT/IN | 0 | 3 |
| LN97 | *Macaca mulatta* | 4 | TNPRC | Male | 4.3 | 2.0 x 10^6^ | IT/IN | 20 | 3 |
| LR09 | *Macaca mulatta* | 4 | TNPRC | Male | 4.8 | 2.0 x 10^6^ | IT/IN | 20 | 3 |
| MD42 | *Macaca mulatta* | 3 | TNPRC | Male | 5.0 | 2.0 x 10^6^ | IT/IN | 20 | 3 |
| MF22 | *Macaca mulatta* | 3 | TNPRC | Male | 5.0 | 2.0 x 10^6^ | IT/IN | 20 | 3 |
| MC12 | *Macaca mulatta* | 3 | TNPRC | Male | 5.3 | 2.0 x 10^6^ | IT/IN | 6 | 3 |
| LR41 | *Macaca mulatta* | 4 | TNPRC | Male | 5.7 | 2.0 x 10^6^ | IT/IN | 6 | 3 |
| ME55 | *Macaca mulatta* | 3 | TNPRC | Male | 5.8 | 2.0 x 10^6^ | IT/IN | 6 | 3 |
| LV40 | *Macaca mulatta* | 4 | TNPRC | Male | 5.6 | 2.0 x 10^6^ | IT/IN | 6 | 3 |
| LM12 | *Macaca mulatta* | 4 | TNPRC | Male | 6.1 | 2.0 x 10^6^ | IT/IN | 2 | 3 |
| LT54 | *Macaca mulatta* | 3 | TNPRC | Male | 6.2 | 2.0 x 10^6^ | IT/IN | 2 | 3 |
| MG10 | *Macaca mulatta* | 3 | TNPRC | Male | 6.0 | 2.0 x 10^6^ | IT/IN | 2 | 3 |
| IR17 | *Macaca mulatta* | 11 | TNPRC | Male | 6.3 | 2.0 x 10^6^ | IT/IN | 2 | 3 |
| LP79 | *Macaca mulatta* | 4 | TNPRC | Male | 5.5 | 2.0 x 10^6^ | IT/IN | 20 | 75 |
| LR93 | *Macaca mulatta* | 4 | TNPRC | Male | 6.0 | 2.0 x 10^6^ | IT/IN | 20 | 75 |
| MC61 | *Macaca mulatta* | 3 | TNPRC | Male | 5.7 | 2.0 x 10^6^ | IT/IN | 20 | 75 |

**Table S2. Nonhuman Primates Used in this Study**

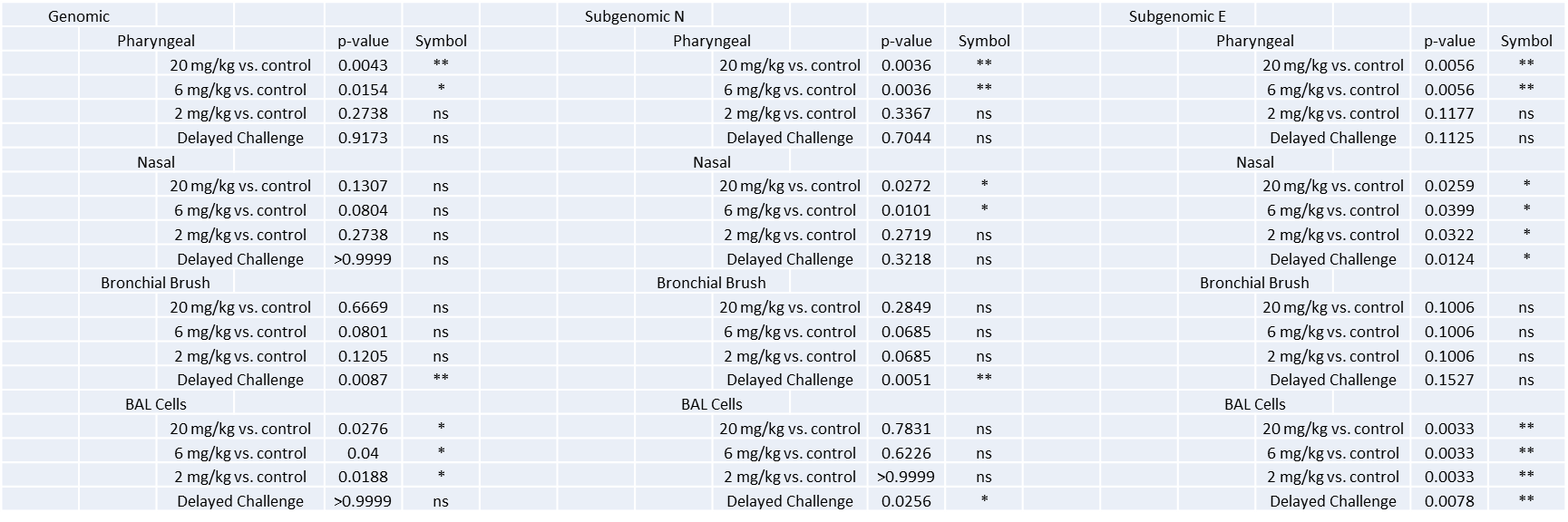

**Table S3. Exact p Values Calculated for RT-qPCR Comparisons.** Groups were compared via Kruskal-Walis test comparing each dosage group to the control group. Asterisks represent significant comparisons (*, p<0.05; **, p<0.01).

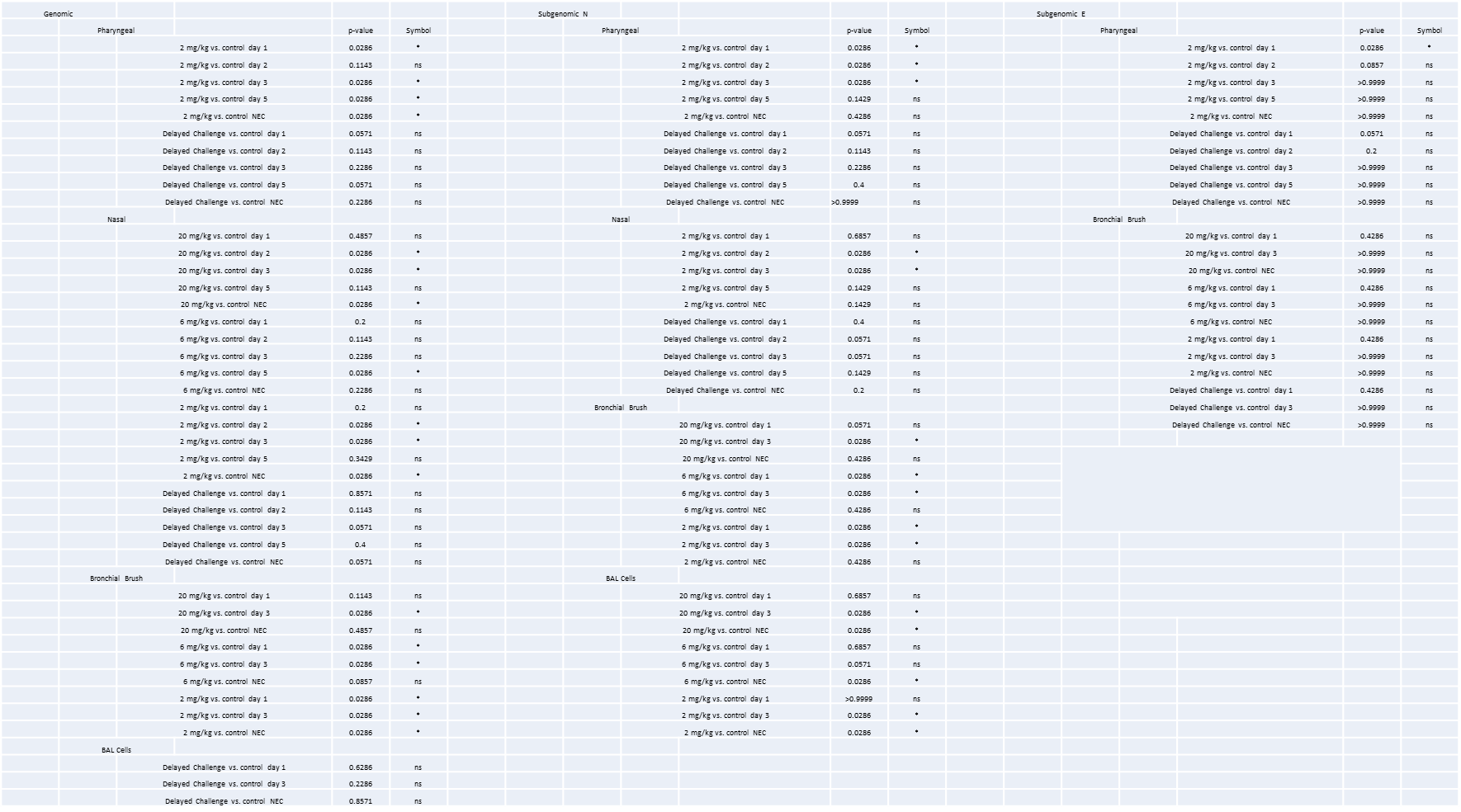

**Table S4. Exact p Values Calculated for Comparison of RT-qPCR Viral Loads Day-by-Day.** Groups were compared via Mann-Whitney test for each comparison not significant overall. Asterisks represent significant comparisons (*, p<0.05; **, p<0.01).

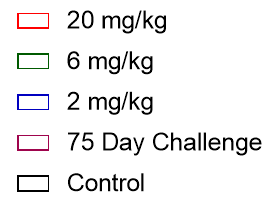

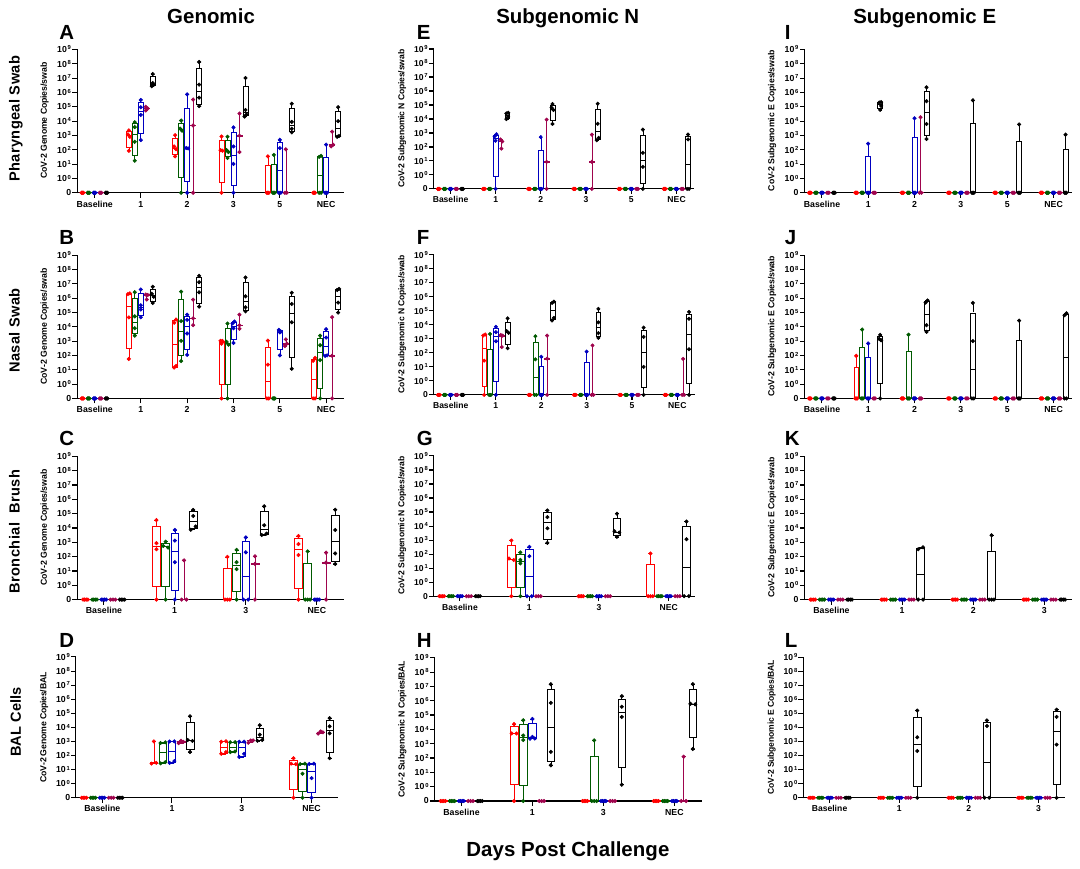

**Figure S1. RT-qPCR Viral Loads over Time.** Viral loads from pharyngeal swabs (A, E, I), nasal swabs (B, F, J), bronchial brushes (C, G, K) and BAL cells (D, H, L) assessed by RT-qPCR for genomic N (A-D), subgenomic N (E-H), and subgenomic E (I-L) content. Data are represented as genome/subgenome copies per swab or BAL per day post challenge.

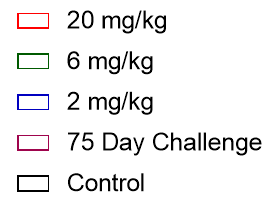

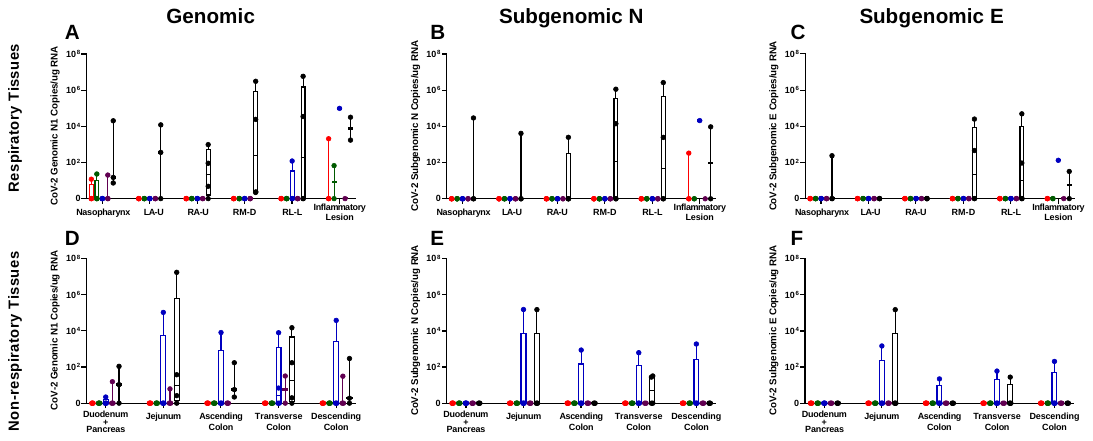

**Figure S2. Viral Loads Found in Tissues at Necropsy.** Viral loads from respiratory (A-C) and non-respiratory (D-F) for genomic (A,D), subgenomic N (B, E) and subgenomic E (C, F) content.

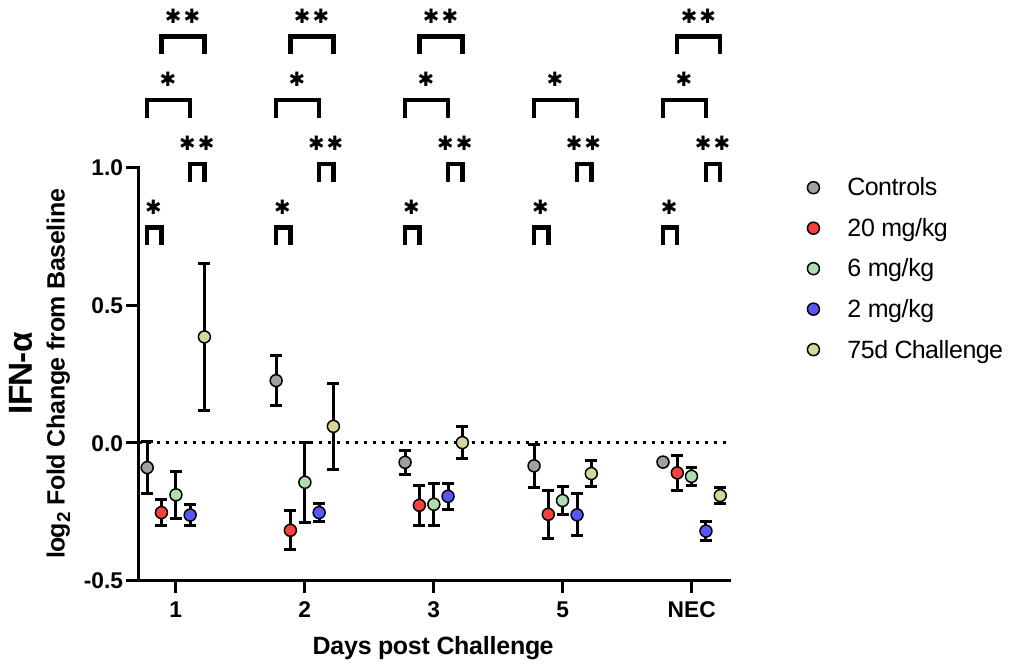

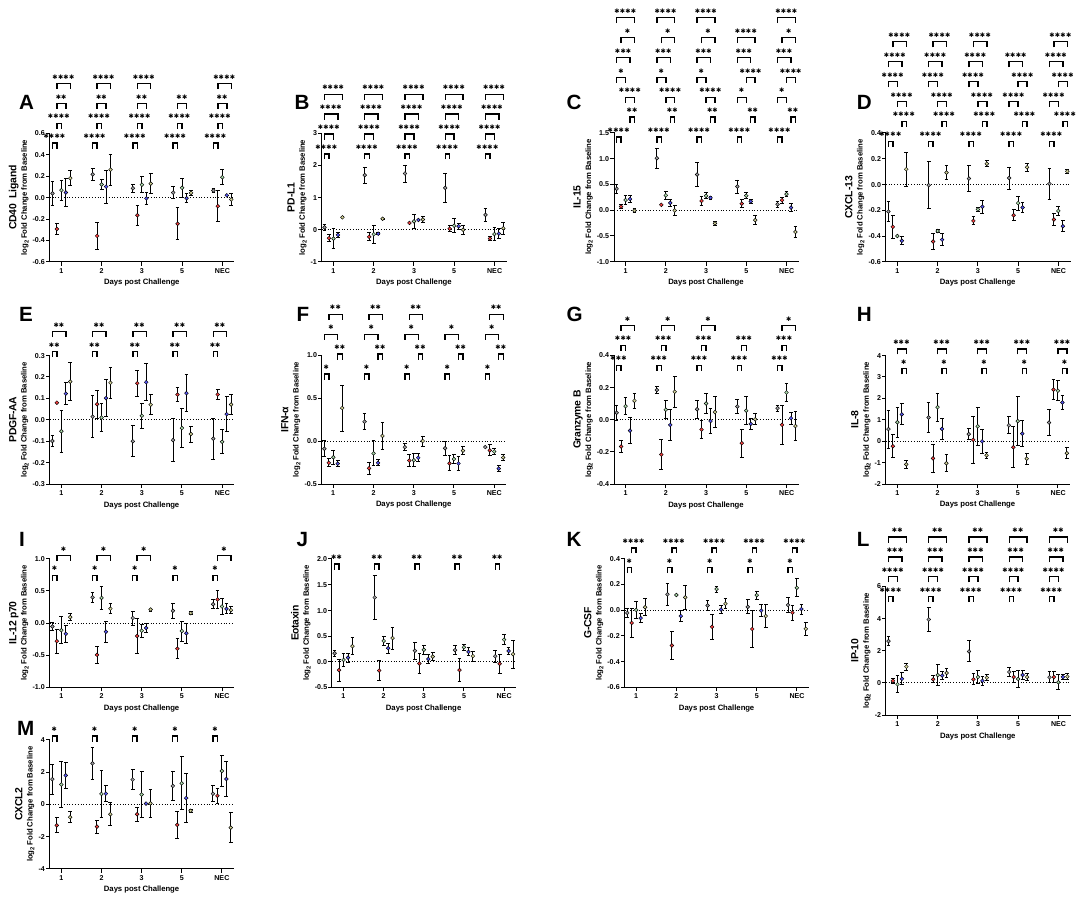

**Figure S3. Additional Cytokine/Chemokine Comparisons.** Representations of log_2_ fold change between dosage groups, with comparisons made via Two-way ANOVA with Tukey’s multiple comparisons test (*, p<0.05; **, p<0.01; ***, p<0.001; ****, p<0.0001).

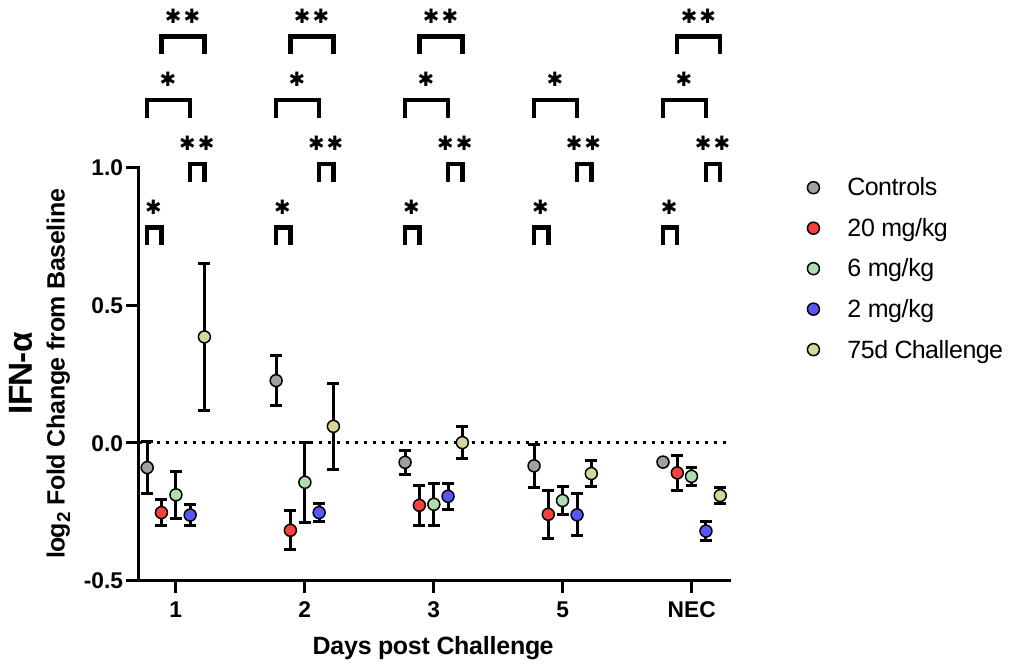

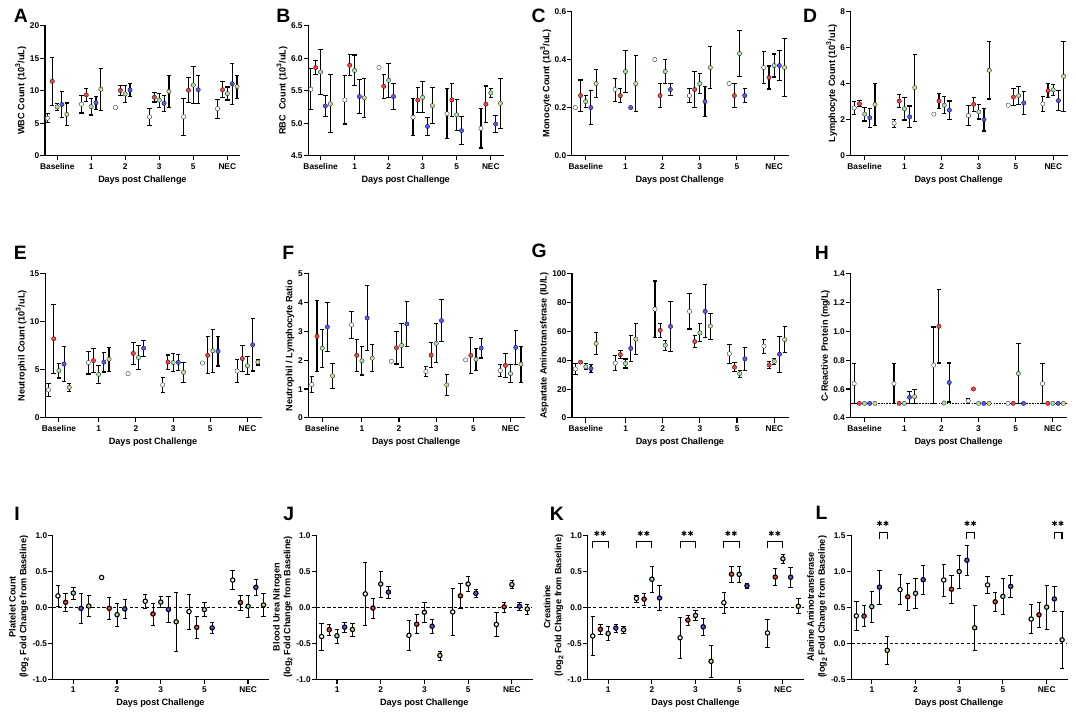

**Figure S4. Hematology and Clinical Chemistries.** Clinical Chemistry and CBC values per dosage group (A-H). Line indicates limit of detection for CRP (H). Log_2_ fold change of selected analyses (I-L), with line indicating no fold change.
